# An Efficient Implicit Tau-Leaping Method for Estimating Second-Order Sensitivities of Stiff Stochastic Biochemical Systems

**DOI:** 10.64898/2026.10.17.759817

**Authors:** Fauzia Jabeen, Mareson Suresh, Silvana Ilie

**Affiliations:** Department of Mathematics, Toronto Metropolitan University Toronto, Ontario, M5B 2K3, Canada

**Keywords:** Stochastic Simulation Algorithm, Stochastic Models of Biochemical Kinetics, Sensitivity Analysis, Implicit Tau-leaping Method, Adaptive Time-stepping

## Abstract

Stochastic modelling has become increasingly important for studying biochemical systems at the cellular level. One of the most commonly used stochastic discrete models of biochemical systems is the Chemical Master Equation. Sensitivity analysis is a fundamental tool for examining how model output changes with variations in model parameters, while second-order sensitivities offer insight into parameter interactions, and provide essential information for parameter estimation and experimental design. Finite-difference approximation of second-order sensitivities can be computationally challenging, especially for stiff biochemical systems, for which reactions occur on widely different scales in time. We propose a finite-difference strategy that uses an adaptive implicit tau-leaping method to estimate second-order parametric sensitivities efficiently. Numerical experiments on several stiff models of practical interest show that the proposed method achieves accuracy comparable to existing approaches while substantially reducing the computational cost.

## 1 Introduction

Biochemical processes in living cells display inherent randomness which arises as a consequence of unpredictable interactions among various molecules. When at least one of the reacting species is present in low copy numbers, as is common for genes, regulatory proteins and signalling molecules, the fluctuations at the molecular level can result in behaviour that the deterministic reaction rate equations cannot capture [9, 17, 33]. Advances in experimental research have led to increasing efforts to develop stochastic models which accurately portray a broad range of biological systems [29, 36].

The Chemical Master Equation (CME) is a widely adopted stochastic discrete model of well-stirred biochemical reaction networks [23, 11]. It models the system state as a Markov process. The direct solution of the Chemical Master Equation is intractable for most realistic biochemical networks. Exact trajectories of the underlying Markov process may be generated by Gillespie’s stochastic simulation algorithm (SSA) [12, 13], and by the more efficient Next Reaction Method [10]. Exact methods are statistically unbiased but they become prohibitively expensive when fast reactions occur in the system. To address this challenge, Gillespie proposed the explicit tau-leaping method [14] which improves computational efficiency by allowing multiple reaction events to occur over a preselected time step *τ*. If the leap condition is satisfied, which means that the propensities remain approximately constant during the step, then the number of firings of each reaction may be approximated by a Poisson distribution. Adaptive step-size selection techniques [6] further improve the efficiency while maintaining the accuracy of the method. This approach has also been extended to stochastic reaction-diffusion systems [30]. Several other numerical strategies have been designed for simulating stochastic discrete models of well-stirred biochemical networks [8, 15, 19, 24, 37, 40, 41] (see also the related references therein).

A large number of biochemical networks exhibit stiff dynamics, with reactions firing on widely separated timescales [5, 7, 15]. For such networks, exact simulation strategies become computationally impractical and the explicit tau-leaping method must take prohibitively small time steps for stability. This problem can be overcome using the implicit tau-leaping [34] to which the effective adaptive explicit-implicit scheme of step selection from [7] is applied. For models involving significantly different timescales, the step sizes of the explicit tau-leaping scheme may be constrained by the fastest reactions, requiring very small time steps to maintain numerical stability. In contrast, the implicit tau-leaping method, in which the drift term is treated implicitly, permits substantially larger time steps while maintaining stability. This makes the method more efficient of systems characterized by stiffness. For networks approaching a steady-state, the implicit tau-leaping strategies can damp stochastic fluctuations, as demonstrated in [34].

Parametric sensitivity analysis quantifies how model outputs change in response to variations in model parameters, and is key for model calibration and experimental design of stochastic biochemical systems [2, 3, 16, 21, 18, 31]. Several first-order sensitivity estimators for the Chemical Master Equation have been developed in the literature, including the coupled finite-difference (CFD) method of [1], the Girsanov likelihood-ratio method [32] and the common reaction path and common random number schemes [35]. In [27], a coupled explicit tau-leaping finite-difference estimator for first-order sensitivities was proposed for moderately stiff systems and it was subsequently extended to a coupled implicit tau-leaping estimator for stiff systems [28].

Estimating second-order sensitivities of the expected outputs with respect to model parameters (the Hessian entries) is an increasingly important problem. However, unlike the first-order sensitivity analysis, the second-order sensitivity analysis for stochastic models is considerably less studied. The elements of the Hessian matrix are useful for Newton-type optimization, for parameter inference and control design, for practical parameter identifiability and the mixed sensitivities help uncover interactions between pairs of parameters, that are not revealed by the first-order sensitivities [18, 21, 42]. Finite-difference approximations of second-order parametric sensitivities in stochastic discrete models present a significant challenge compared to their first-order counterparts, as they can amplify the effects of the simulation noise. Therefore, accurate approximations require strong coupling techniques together with reliable Monte Carlo methods. The CFD technique [1] was extended to second-order derivatives (CFD-2) in [42], utilizing Kurtz’s random time change representation and a modified next reaction method. The presence of widely separated timescales, which happens often in biochemical networks arising in applications, can make strategies based on exact stochastic simulation techniques such as the CFD-2 prohibitively expensive. In [20], the coupled tau-leaping method of second order (CTL-2) was proposed, which combined second-order coupled finite-difference schemes with variable step size explicit tau-leaping strategies. This method is appropriate for moderately stiff models of biochemical networks. Still, since it is an explicit technique, it becomes inefficient for very stiff models.

In this paper, we address this challenge by developing a finite-difference method to estimate second-order sensitivities based on the implicit tau-leaping scheme. The proposed method is called the Coupled Implicit-Tau of second-order (CIT-2) strategy. For each Hessian entry, four sample paths, one for the nominal, two for the singly perturbed and one for the doubly-perturbed systems are simulated. We employ the same relative perturbation parameter, *h*, thus to each kinetic parameter *c*_*j*_ it corresponds an absolute perturbation *h*_*j*_ = *h c*_*j*_. To ensure a significant reduction of the finite-difference estimator variance, the four processes are strongly coupled, which is essential as second-order differencing magnifies the simulation noise to a much greater extent than in the case of estimating sensitivities of first-order by finite-difference schemes. By generating the sample paths with the implicit rather than the explicit tau-leaping method, the estimator remains stable and effective when applied on stiff to very stiff systems. Our technique adapts a variable time-stepping strategies for implicit tau-leaping proposed in [7] and extends the coupled implicit tau first-order (CIT) sensitivity estimator of [28], to second-order while ensuring a strong coupling of the four processes is applied. This coupling leads to a reduction in the computational cost of the estimation. Our approach draws on ideas from the coupled finite-difference framework introduced in [1], but it is aimed at a wide class of stiff, realistic biochemical systems, for which finite-difference methods relying on exact stochastic simulation techniques are computationally demanding.

The remainder of this paper is organized as follows. Section 2 reviews the background on the Chemical Master Equation model, exact stochastic simulation strategies and the explicit and implicit tau-leaping schemes. Section 3 discusses the coupled finite-difference framework for sensitivity estimation, existing second-order sensitivity estimators and proposes a new implicit coupled tau-leaping estimator for second-order sensitivity analysis. In Section 4, we present the advantages of the new sensitivity estimator compared to existing ones, by testing it on several stiff biochemical systems of practical interest. Section 5 concludes with a discussion and directions for future work.

## 2 Background

### Stochastic Discrete Biochemical Networks

Consider a well-mixed biochemical system for which *N* molecular species *S*_1_, …, *S*_*N*_ interact through *M* reaction channels *R*_1_, …, *R*_*M*_ inside a fixed volume which is kept at constant temperature. The state of the system at time *t* is the vector of molecular populations **X**(*t*) = (*X*_1_(*t*), …, *X*_*N*_ (*t*))^*T*^, such that *X*_*i*_(*t*) is an integer denoting the molecular abundance of species *S*_*i*_ at time *t*. The initial state of the system is specified, **X**(*t*_0_) = **x**_0_.

Each reaction channel *R*_*ℓ*_ is associated with two quantities. The first is its stoichiometric vector *ν*_*ℓ*_ = (*ν*_1l_, *ν*_2l_, …, *ν*_*Nl*_)^*T*^, where *ν*_*il*_ is the change in the population of *S*_*i*_ produced by one firing of *R*_*ℓ*_, so that after one reaction takes place the system state **x** is updated to **x**+***ν***_***ℓ***_. The second is its propensity function *a*_*ℓ*_, defined by *a*_*ℓ*_(**x**) *dt* = the probability that one *R*_*ℓ*_ reaction occurs in [*t, t* + *dt*), subject to **X**(*t*) = **x**. Under mass-action kinetics with stochastic rate constant *c*_*ℓ*_, the propensity of a zero-order reaction is *a*_*ℓ*_ (**x**) = *c*_*ℓ*_, of the first-order reaction *S*_*i*_→ products, *a* (**x**) = *c x*_*i*_, of the bimolecular reaction event *S*_*i*_ + *S*_*k*_→ products with *i* ≠ *k, a*_*ℓ*_(**x**) = *c*_*ℓ*_*x*_*i*_*x*_*k*_, and of the dimerization *S*_*i*_ + *S*_*i*_ →products, *a*_*ℓ*_(**x**) = *c*_*ℓ*_*x*_*i*_(*x*_*i*_ −1)*/*2, which equals the number of distinct pairs of *S*_*i*_ molecules.

The system state **X**(*t*) is represented by a continuous-time, discrete-state Markov process. Its probability distribution, defined by *P* (**x**, *t* |**x**_0_, *t*_0_) is the probability that **X**(*t*) = **x**, given that **X**(*t*_0_) = **x**_0_, satisfies the Chemical Master Equation [11]

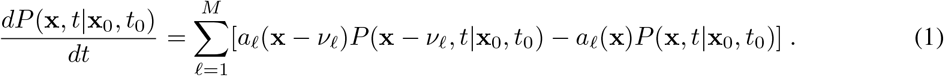

The system state depicted by the Markov process **X**(*t*) also satisfies the random time change representation (RTC) [23, 22],

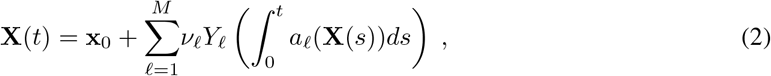

where *Y*_*ℓ*_ for 1 ≤*ℓ* ≤ *M* are independent Poisson processes of unit rate and the initial state is **X**(*t*_0_) = **x**_0_.

Note that 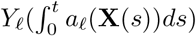 counts the number of reactions *R*_*ℓ*_ that happened within [*t*_*0*_, *t*].

Exact realizations of the Markov process governed by (1) can be computed using Monte Carlo simulation methods, such as Gillespie’s Stochastic Simulation Algorithm [12, 13] or the next reaction method [10]. However, in systems exhibiting stiffness, such methods are computationally intensive. Indeed, as exact strategies simulate each reaction event individually, their computational cost can become considerable for networks with fast reactions.

### Tau-leaping methods

The tau-leaping strategy due to Gillespie [14] accelerates the simulation by advancing the system over a preselected time step *τ* which leaps over many reactions. The time-step must satisfy the leap condition: *τ >* 0 is small enough such that no propensity function has a significant variation during [*t, t* + *τ*). When this assumption is valid, that is the propensities are almost constant over the step, the number of reaction events *R*_*ℓ*_ in [*t, t* + *τ*) may be estimated by a Poisson random variable *P*_*ℓ*_*(a*_*ℓ*_(**x**) *τ*, of mean and variance *a*_*ℓ*_ (**x**)*τ*. Thus, we obtain the *explicit tau-leaping* strategy,

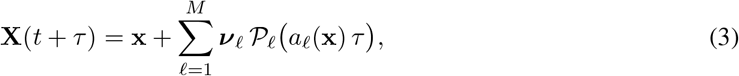

if *X*(*t*) = **x**. Here {*P*_*ℓ*_ (*a*_*ℓ*_ (**x**) *τ*_*ℓ*=1,…,*M*_ are independent Poisson random variables. The explicit tau-leaping method (3) was introduced by Gillespie [14].

Numerous realistic biochemical networks evolve on multiple time-scales and are therefore stiff: fast reaction events, often reversible pairs that reach equilibrium, along with much slower reactions are present in the system. For such networks, the explicit tau-leaping scheme will take very small time steps determined by the fastest reactions to maintain numerical stability, and thus it becomes computationally intensive. The implicit tau-leaping technique proposed in [34] overcomes this difficulty by considering an implicit term in the drift but not in the noise. Specifically, the *implicit tau-leaping* scheme [34] is given by

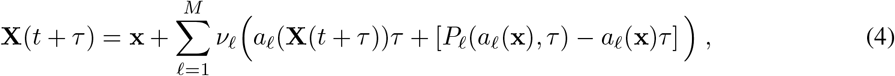

provided that *X*(*t*) = **x**. This strategy has better stability properties than the explicit tau-leaping scheme (3), as its step size is specified by the slow dynamics rather than the fastest one [34]. At each step, the nonlinear system (4) requires an implicit solver, such as Newton’s method, to approximate the system state **X**(*t* + *τ*).

### Variable time-stepping for the implicit tau-leaping method

A variable time-stepping technique for (4) was designed by Cao et al. [7]. This technique employs a variant of the leap condition and determines the leap length based on the reactions which are not in partial equilibrium, after identifying the fast reversible channels. Also, the variable time-stepping method handles critical reactions separately in order to avoid spurious negative population numbers.

The efficiency of the time-stepping selection strategy for the implicit tau-leaping scheme is based on the observation that the dynamics of the system is mainly decided by the reactions which are not in partial equilibrium. Consequently, the reactions that are in partial equilibrium may be ignored when selecting the step length. The strategy identifies the reversible reactions that have reached partial equilibrium. These are often fast and excluding them when selecting the next step can lead to significant efficiency gains when simulating stiff systems. A reversible pair of reaction channels (*R*_+_, *R*_−_), with propensities *a*_+_(**x**) and *a*_−_(**x**), is considered to be in *partial equilibrium* provided that the following condition is satisfied

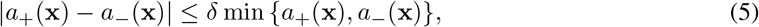

for some small parameter *δ >* 0. The set of reactions which are not in partial equilibrium are represented by *J*_*ne*_.

The tau-leaping strategy may fire a reaction more times than its reactant amounts allow and this will lead to physically unrealistic negative population numbers. This issue is avoided by partitioning all reactions into critical and non-critical. For a reaction *R*_*ℓ*_ with propensity *a*_*ℓ*_ (**x**) *>* 0, we denote by

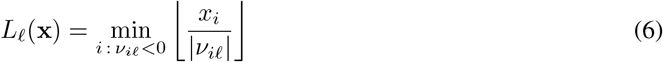

the maximum number of times a reaction *R*_*ℓ*_ may occur before depleting one of its reactants. The minimum in (6) is taken over the reactants that *R*_*ℓ*_ consumes. We used the notation ⌊ · ⌋for the greatest integer less than the input value. The reaction is considered *critical* when *L*_*ℓ*_(**x**) *< n*_*c*_, where *n*_*c*_ is a threshold parameter, typically taking values between 2 and 20. Critical reactions are simulated according to a stochastic simulation algorithm applied to critical reactions. They are generated one at a time, so that at most one critical reaction event may happen over a step. By contrast, the non-critical reactions are computed according to the implicit tau-leaping strategy. We denote by *J*_*cr*_ and *J*_*ncr*_ the set of critical and non-critical reactions, respectively, and by *I*_*rs*_ and *I*_*ncr*_ the set of indices of all reactant species and of those for non-critical reactions, respectively. Finally, the set of non-critical and not in partial equilibrium reactions are indicated by *J*_*necr*_ = *J*_*ne*_ ∩ *J*_*ncr*_.

#### Stepsize for critical reactions

The critical reactions are advanced one at a time, using a variant of the stochastic simulation algorithm restricted to the set of critical reactions, *J*_*cr*_. We make the notation 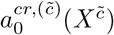 to be the summation of all propensities of critical reactions for the trajectory 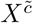 corresponding to the parameter 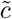. For the critical reactions, we apply the direct method of Gillespie. We generate two samples 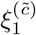 and 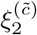 of *U* (0, 1), the uniform random variable on the interval [0, 1]. Then, the index *j*_*cr*_ of the next critical reaction is computed according to

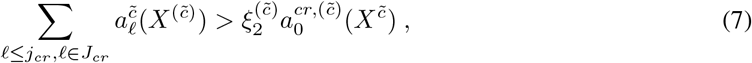

while the time to the next critical reaction event is

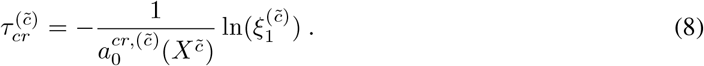

#### Stepsize for not in partial equilibrium, non-critical reactions

Following Cao et al. [7], for the reactions which are not critical and also not in partial equilibrium, we impose the leap criteria that the mean and variance of each reactant molecular amount *X*_*i*_ obeys

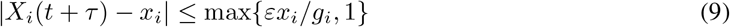

for any *i* ∈ *I*_*rs*_, where 0 *< ε* ≪1 is the tolerance for time-adaptivity. The factor *g*_*i*_ is given by

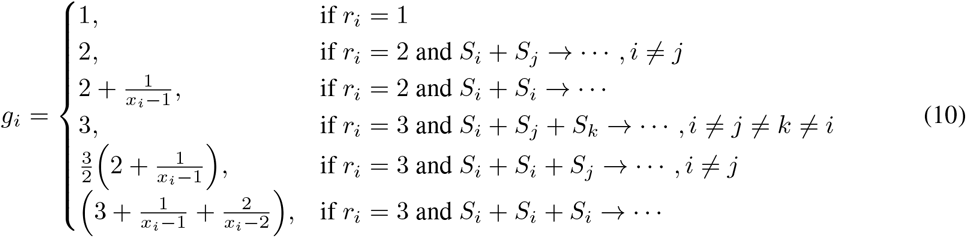

where *r*_*i*_ to be the highest order of a reaction in which the species *S*_*i*_ is a reactant (see [6] for details). Approximating the change in *X*_*i*_ over the step by its mean and variance, computed over the reactions in *J*_*necr*_ only, and imposing (9) leads to the following stepsize *τ* for the implicit tau-leaping strategy

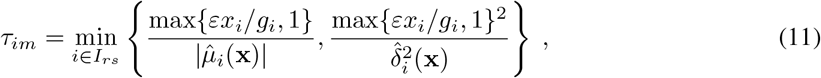

with the auxiliary quantities

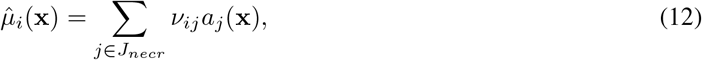

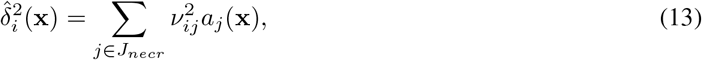

for *i* ∈*I*_*rs*_.

The proposed method for second-order sensitivity estimation for the Chemical Master Equation is implemented with a variation of the *τ* -selection technique of Cao et al. [7].

## 3 Second-Order Parametric Sensitivity

Sensitivity analysis quantifies how parameter variations affect the dynamics of biochemical networks, assisting in identifying key parameters, assessing model robustness, and guiding experimental design and parameter estimation. Local sensitivity analysis examines the effects of small variations in model parameters, such as initial molecular counts and reaction rate parameters, on the system dynamics. High sensitivity for one parameter suggests that accurate estimation of that parameter value is critical for correct predictions, whereas low sensitivity shows that small variations in the value of that parameter have minor impact on the system behaviour.

Assume that the biochemical model depends on the vector of *M* parameters *c* = (*c*_1_, …, *c*_*M*_)^*T*^ and denote by **X**(*t, c*) the state of the system at time *t* corresponding to the parameter values *c*. If **f** is a smooth function of the system state, and if E[·] denotes the expected value, then the second-order sensitivity with respect to parameters *c*_*j*_ and *c*_*k*_ is

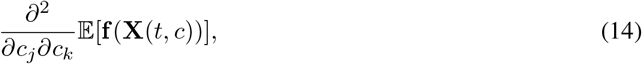

for *j, k* ∈{1, …, *M*}. The second-order sensitivities (14) are the entries of the Hessian of the expected value of the function of the state **X**(*t, c*). In this paper, the parameters of interest are the kinetic parameters *c*_*j*_, corresponding to reactions *R*_*j*_, for *j* ∈{1, …, *M*}. Finite-difference techniques may be utilized to estimate second-order sensitivities for stochastic discrete models of biochemical networks. Let *e*_*j*_ be the unit vector in ℝ ^*M*^ with the *j*-th entry being 1. A forward finite-difference estimation of sensitivity (14) can be expressed as

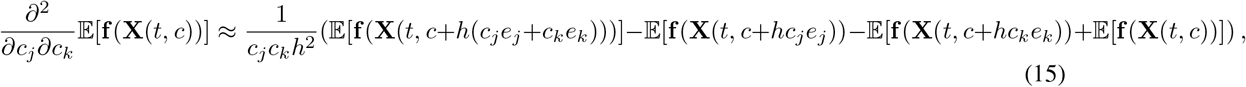

with 0 *< h <* 1 being a relative perturbation, and *c*_*j*_ and *c*_*k*_ the selected parameters. Such a method involves the nominal process **X**(*t, c*), the processes perturbed in each parameter, **X**(*t, c*+*hc*_*j*_*e*_*j*_) and **X**(*t, c*+*hc*_*k*_*e*_*k*_), and the process perturbed in both parameters, **X**(*t, c* + *h*(*c*_*j*_*e*_*j*_ + *c*_*k*_*e*_*k*_)).

The expectations in (15) are estimated using Monte Carlo methods. If *X*(*t, c*) represents a sample path of the stochastic process **X**(*t, c*), a realization of the finite-difference scheme (15) is represented by

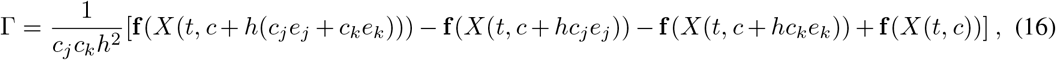

and the sample mean of such realizations is an estimation of (14). Reducing the number of realizations necessary for a certain accuracy can be done by lowering the variance of the sensitivity estimator. This leads to a reduced computational cost for approximating the parametric sensitivity with the desired accuracy. Lowering the variance of the estimator can be achieved by coupling the four processes.

The CFD-2 strategy [42] extends to second-order parametric sensitivity approximation the coupled finite-difference method for first-order sensitivities of Anderson [1]. This strategy proposed a strong coupling of the four processes [42]

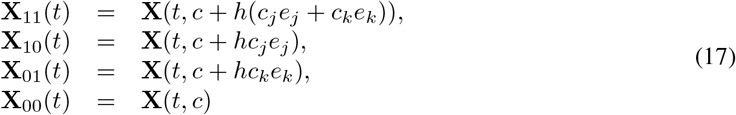

and used a variation of the Next Reaction Method to generate them. The quadruple processes (17) can be expressed in a unified form as

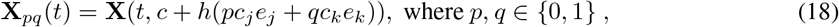

and they satisfy the initial condition **X**(0, *c* + *h*(*pc*_*j*_*e*_*j*_ + *qc*_*k*_*e*_*k*_)) = **x**_0_. Below, we shall describe briefly the coupling introduced in [42]. For each reaction channel *R*_*ℓ*_, 1≤*ℓ* ≤*M*, we represent its propensity for each of the four process by

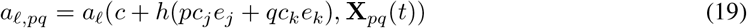

for *p, q* ∈{0, 1} where, for simplicity, the dependence on time and on the parameters is omitted.

As described in [42], each counting process *Y*_*ℓ*_ of the random time change representation (2) is decomposed into subprocesses shared among the four continuous in time discrete in space Markov processes (18). Each subprocess is indexed by the binary sequence *b* = *b*_1_*b*_2_*b*_3_*b*_4_ such that *b*_*i*_ ∈{0, 1}. We define the set of all these binary sequences by

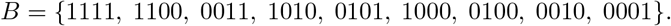

and the subsets relevant for each process by

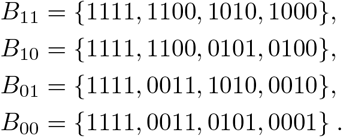

Also, we denote the intensities used for coupling by

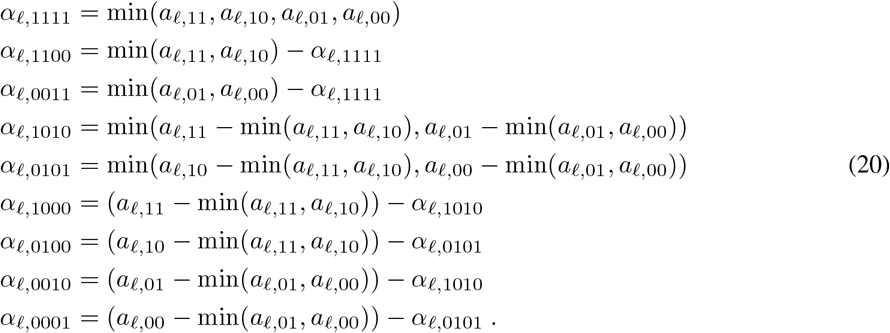

where *α*_*ℓ,b*_ are such that 1 ≤*ℓ* ≤ *M* and *b* ∈*B*.

To obtain a tight coupling of the four processes, the CFD-2 strategy [42] uses the random time change representation (2), and expresses these processes as

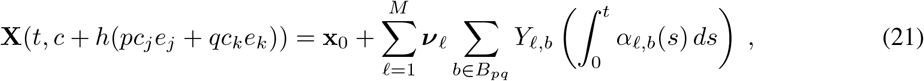

where *p, q* ∈ {0, 1}. Furthermore, note that 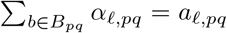 for every reaction *R*_*ℓ*_, with 1 ≤*ℓ* ≤*M*, and every process. Consequently, each of the correlated processes above is identically distributed to its corresponding uncoupled process.

### 3.1 Coupled Implicit-Tau Strategy for Estimating Second-Order Sensitivities

Below, we develop an implicit tau-leaping method for the efficient and accurate estimation of second-order sensitivities in stochastic discrete models of well-stirred biochemical systems. The method utilizes finite-difference sensitivity estimation and an implicit tau-leaping scheme. The algorithm uses an adaptive time step selection, which allows larger leaps while maintaining accuracy and numerical stability. This provides an effective strategy for approximating second-order sensitivities for biochemical network models with multiple timescales and it represents a significant extension of the existing finite-difference techniques.

The implicit *τ* -leap strategy is particularly advantageous for stiff to very stiff biochemical reaction networks, which exhibit both slow and fast dynamics. The presence of fast and slow reactions is common in biochemical reaction networks arising in applications. For such networks, the exact stochastic simulation techniques can be computationally demanding since they simulate each reaction, one at a time. The CFD-2 technique [42] employs an exact algorithm to generate the four sample paths, simulating every reaction on each path, and it is well suited for non-stiff models of biochemically reacting systems. However, when the system contains fast reactions, this computational cost becomes prohibitive. The CTL-2 method [20] reduces the cost by considering approximate explicit tau-leaping techniques, which skip over many reactions and it is efficient on models which are moderately stiff. Nonetheless, for very stiff systems, this method require small time steps for maintaining numerical stability, thus it becomes inefficient on such models. By contrast, the proposed implicit *τ* -leaping method, CIT-2, permits larger time steps while incorporating the effects of fast reactions; this can greatly reduce the computational cost of sensitivity estimation with insignificant loss in accuracy.

The coupling of the quadruple processes (17) in the CIT-2 method for estimating second-order sensitivities with respect to parameters *c*_*j*_ and *c*_*k*_ is constructed as follows:

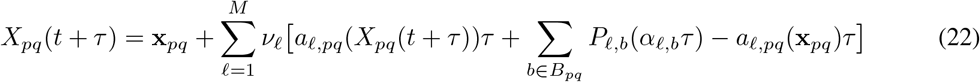

where *X*_*pq*_(*t*) = **x**_*pq*_ for *p, q* ∈ {0, 1} and *P*_*ℓ,b*_(*α*_*ℓ,b*_*τ*) are independent Poisson random variables of mean and variance *α*_*ℓ,b*_*τ*, for 1 ≤*ℓ* ≤ *M* and *b* ∈*B*. For *p, q* ∈ {0, 1} and 1 ≤*ℓ* ≤ *M*, we denoted by

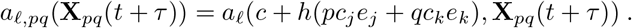

The proposed coupled implicit *τ* -leaping (CIT-2) method correlates the nominal and three perturbed processes using shared Poisson random variables and uses finite-difference approximations to estimate second-order sensitivities for stochastic discrete biochemical models. These four coupled processes are expected to be strongly correlated, which leads to a reduction in the variance of the estimation. The CIT-2 algorithm employs a variable time-stepping strategy for the implicit tau-leaping technique, which is adapted from the method by Cao et al. [7]. It generates a sample path for each of the four processes, as required for the finite-difference estimation, with the implicit *τ* -leap scheme on a common sequence of time-steps. Critical reactions are generated one at a time, to avoid negative population numbers, whereas the non-critical reactions are simulated with the implicit tau-leaping scheme. For a derivative of second-order with respect a parameter (*j* = *k*), the two perturbed processes corresponding to *c* + *hc*_*j*_*e*_*j*_ and *c* + *hc*_*k*_*e*_*k*_, share the same parameter values, while the doubly perturbed process has parameter *c* + 2*hc*_*j*_*e*_*j*_. Two distinct paths are simulated for the processes corresponding to *c* + *hc*_*j*_*e*_*j*_ and *c* + *hc*_*k*_*e*_*k*_, to ensure the correct coupling.

Below, we present the proposed CIT-2 algorithm for estimating second-order sensitivities for stochastic discrete models of biochemical networks.

#### Coupled Implicit Tau -2 Algorithm

1. **Input**: final time *T*, adaptivity tolerance *ε*, Netwon solver tolerance *tol*, partial equilibrium tolerance *δ*, critical threshold *n*_*c*_, relative perturbation *h*, indexes *j, k*, initial condition **x**_0_.
2. **Initialization** *t* = 0, *X*_*pq*_ = **x**_0_, for all *p, q* ∈ {0, 1}.
3. **While** *t < T* **do** steps (a) – (j):
  a. **Evaluate the propensities for each path** from (19): for *ℓ*= 1, …, *M* and *p, q* ∈ {0, 1},

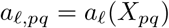
  b. **Identify reactions in partial equilibrium**: for each set of reversible reactions, on each path, verify if they are in partial equilibrium using (5)
  c. **Identify critical reactions for each path**: for *p, q* ∈{0, 1}find the set of reactions not in partial equilibrium *R*_*ℓ*_, with *a*_*ℓ,pq*_ *>* 0, and take

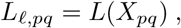

where *L*_*ℓ,pq*_ defined by (6). **Set** *J*_*ncr*_ = {*ℓ* |*L*_*ℓ,pq*_ ≥*n*_*c*_ for any *p, q* ∈{0, 1}}and *J*_*necr*_ the set of non-critical, not in partial equilibrium reactions.
  d. **Compute leap sizes**, *τ*_*im,pq*_, **for noncritical, non partial equilibrium reactions**; for all *p, q* ∈ {0, 1}:
    i. If *J*_*necr*_ = ∅, then *τ*_*ncr,pq*_ = ∞.
    ii. If *J*_*necr*_ = ∅, then find set of indices of reactant species of non-critical reactions *I*_*ncr*_.
    iii. For each *i* ∈*I*_*ncr*_ and each trajectory, find the highest order of reaction *r*_*i*_ and the function *g*_*i*_ defined by (10).
    iv. Calculate the auxiliary values 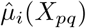 and 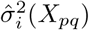 from (12)-(13) and, from (11),

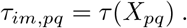
  e. **Compute step sizes**, *τ*_*cr,pq*_, **for critical reactions**; for all *p, q* ∈ {0, 1}:
    i. Find 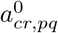 as the sum of the critical reaction propensities for the path *X*_*pq*_.
    ii. Sample *η*_*pq*_ ∼ *U* (0, 1) and generate the time of first critical reaction:

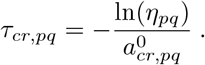
  f. **Find** *τ*, **the step size**: *τ*_*im*_ = min_*p,q*∈{0,1}_{*τ*_*im,pq*_}, *τ*_*cr*_ = min_*p,q*∈{0,1}_{*τ*_*cr,pq*_}, *τ* = min{*τ*_*im*_, *τ*_*cr*_}.
  g. **Determine the number of critical** *R*_*ℓ*_ **firings**, *k*_*ℓ*_ :
    i. If *R*_*ℓ*_ is a critical reaction, then *k*_*ℓ,pq*_ = 0, for any *p, q* ∈ {0, 1}.
    ii. If *τ*_*cr*_ ≤ *τ*_*im*_, then at least one critical reaction fires. For each *p, q* with *τ*_*cr,pq*_ = *τ*_*cr*_, generate *ℓ*^∗^ as a sample of the integer random variable with probability 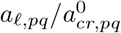, with *ℓ* ∈*J*_*cr*_, 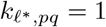.
  h. **Apply implicit tau-leaping over non-critical reactions**:
    i. For *ℓ* ∈*J*_*ncr*_, *b* ∈*B*, calculate *α*_*ℓ,b*_ from (20) and draw the Poisson random variables *k*_*ℓ,b*_ = Poisson(*α*_*ℓ,b*_*τ*)
    ii. Set 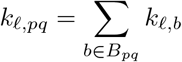.
    iii. Apply Newton’s method for implicit tau-leaping to solve for *Y*_*pq*_ for each *p, q* ∈ {0, 1}:

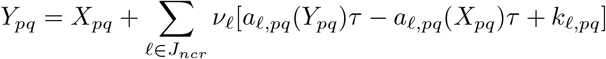
  i. **Advance the time and states on all paths**:
    i. *t* = *t* + *τ*.
    ii. 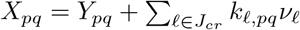, for all *p, q* ∈ {0, 1}.
  j. **Approximate the second order sensitivity**:

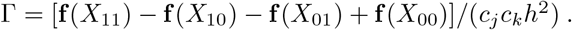

## 4 Numerical Experiments

In this section, we test the proposed CIT-2 technique, on three benchmark models of stiff biochemical systems: the decay-dimerization, a genetic positive feedback loop network and a mutually inhibiting enzymes model. For each system, we generate an ensemble of Monte Carlo trajectories of the same size, using both our new implicit tau-leaping method for second-order sensitivity approximation, CIT-2, and the existing coupled finite-difference scheme, CFD-2. We estimated second-order parametric sensitivities of the expected value of the system state using a forward finite-difference scheme with a relative perturbation parameter *h*. When approximating second-order sensitivities with respect to the rate parameters *c*_*j*_ and *c*_*k*_, the relative perturbation parameter *h* is chosen so that *h* ∈ [0, 1] and the corresponding absolute perturbations satisfy *hc*_*j*_ *<* 1 and *hc*_*k*_ *<* 1. In our simulations, we set the critical threshold value to *n*_*c*_ = 10. We compared the accuracy, the standard deviation of the sensitivity estimation and computational cost of the two strategies, as follows. For both methods, we plotted the time evolution of the means and standard deviations of the trajectories generated at the nominal values of the parameters. Furthermore, we computed the estimated second-order parametric sensitivities and the standard deviations of the sensitivity estimators. Finally, we calculated the speed-ups for the CIT-2 over the CFD-2 for each model, by using the formula

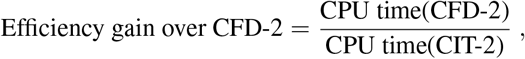

and we found a significant improvement in the computational cost of our method compared to the existing one, while preserving the accuracy, for a range of stiff to very stiff models of stochastic biochemical models, of small to moderate sizes. For the first model, we also examine the dependence of the CIT-2 and CFD-2 sensitivity estimator variances on the relative parameter *h*.

### 4.1 Decay-Dimerization Model

The decay-dimerization model [7, 14] consists of four reactions and three species. A monomer *S*_1_ decays, two monomers bind reversibly to form a dimer *S*_2_, and *S*_2_ can be transformed into a third species, *S*_3_. These species are subject to the reactions shown in Table 1. Table 1 also lists the reaction propensities and the corresponding rate parameter values. The model is investigated on the time interval [0, 1], with initial conditions [*X*_1_(0), *X*_2_(0), *X*_3_(0)] = [400, 800, 0]. In this setting, the decay-dimerization model exhibits multiple time-scales, with the reversible reactions being much faster than the others.

**Table 1:** Decay-dimerization model.

|  | Reaction | Propensity | Parameter value |
| --- | --- | --- | --- |
| $R_1$ | $S_1 \xrightarrow{c_1} \emptyset$ | $a_1 = c_1 X_1$ | 0.05 |
| $R_2$ | $S_1 + S_1 \xrightarrow{c_2} S_2$ | $a_2 = c_2 \frac{X_1(X_1-1)}{2}$ | 50 |
| $R_3$ | $S_2 \xrightarrow{c_3} S_1 + S_1$ | $a_3 = c_3 X_2$ | $10^6$ |
| $R_4$ | $S_2 \xrightarrow{c_4} S_3$ | $a_4 = c_4 X_2$ | 0.05 |

An ensemble of 100, 000 quadruples of trajectories was generated for this model. For the CIT-2 method, we selected the tolerance *ε* = 0.001. In Figure 1(a), we display the mean of the number of *S*_2_ molecules over time, simulated for the nominal values of the parameters, with both the adaptive implicit *τ* –leaping method and the exact next reaction method (NRM). These plots show an excellent agreement for the means produced by the two methods, for the chosen tolerance *ε*. The dependence of the standard deviation of the *S*_2_ molecular amounts on time with each of these methods is presented in Figure 1(b).

**Figure 1:**
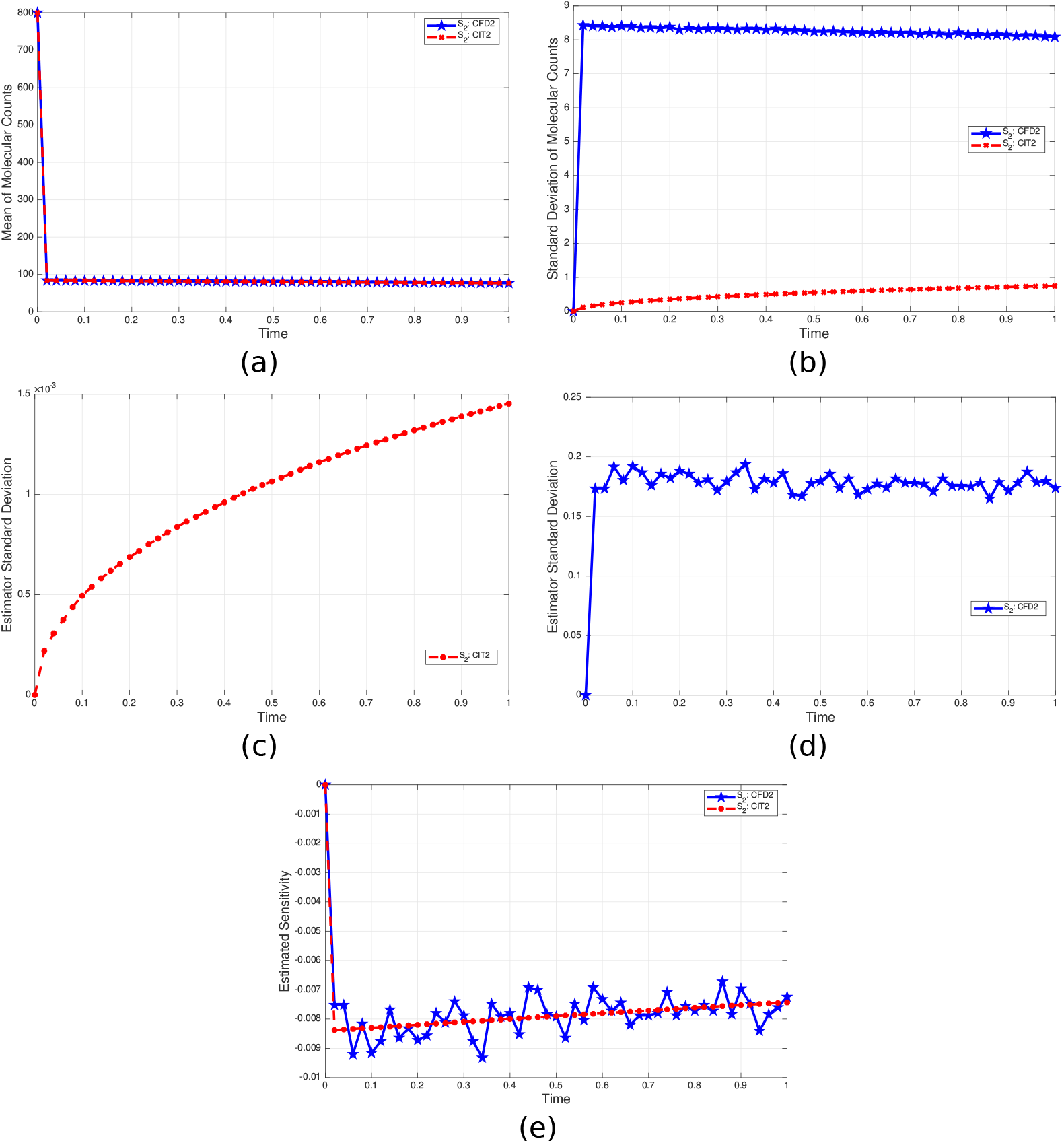
Decay-dimerization model: (a) Mean, (b) Standard deviation of the molecular numbers of *S*_2_ using the implicit tau-leaping and the next reaction methods for the nominal values of the parameters. (c) Standard deviation of the CIT-2 and (d) Standard deviation of the CFD-2 sensitivity estimator, both for the molecular numbers of *S*_2_ with respect to *c*_2_, (e) Second-order sensitivity estimations as above with the CFD-2 and CIT-2 schemes. For each method, we computed 100,000 quadruples of trajectories.

We estimated the *second-order sensitivities with respect to parameter c*_2_ with the CIT-2 and CFD-2 algorithms, for a relative perturbation *h* = 0.5 which is 1% of the value of the perturbed parameter *c*_2_. Figure 1(c) and Figure 1(d) depict the temporal evolution of the variance of the CIT-2 and CFD-2 sensitivity estimators for species *S*_2_, respectively. By comparing the two graphs, it can be observed that the CFD-2 estimator produces a significantly larger variance compared to the CIT-2 estimator, a difference of two orders of magnitude. These findings are confirmed by Figure 1(e), which depicts how the CIT-2 and CFD-2 estimates of the sensitivity 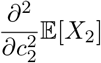 change over time; the CIT-2 approximation is significantly smoother than that computed with the CFD-2 scheme.

The dependence of the sensitivity estimator variance on the perturbation size for each of the two finite-difference methods was examined experimentally. Figure 2 displays the log of the estimator variances versus the log of the perturbation *h*. The results suggest a CIT-2 variance of order O (*h*^−1^) and a CFD-2 variance of order O (*h*^−2^), for this model.

**Figure 2:**
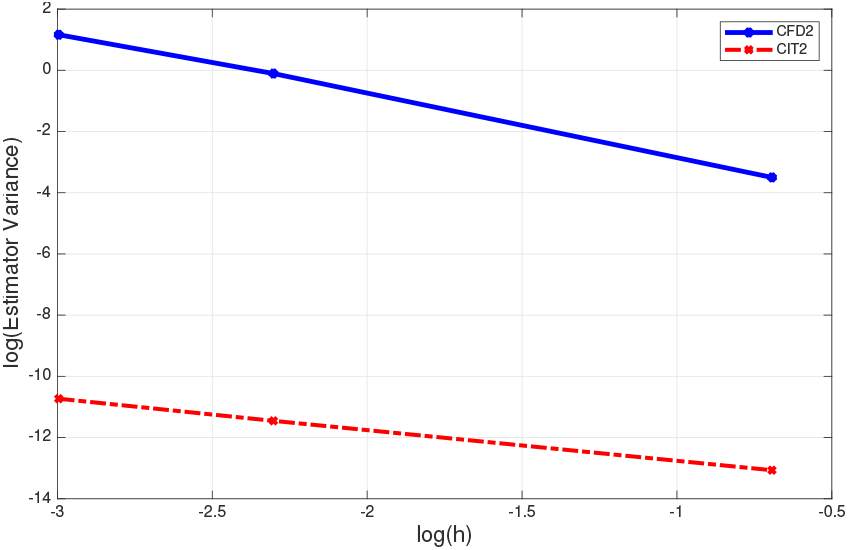
Decay-dimerization model: log of variance of the sensitivity estimator of the *S*_2_ molecular amounts versus log(*h*), for the CIT-2 and CFD-2 algorithms for the double derivative with respect to *c*_2_.

The speed-up of the CIT-2 over the CFD-2 algorithm was computed and found to be

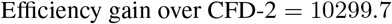

This confirms that the CIT-2 algorithm *significantly outperforms* CFD-2 in terms of computational efficiency on this stiff model, while being very accurate.

### 4.2 Genetic Positive Feedback Loop Model

The second model is the genetic positive feedback loop system introduced by Bennett et al. [4]. The model describes a single gene transcribing a single protein. It comprises five biochemical species: the protein monomer *S*_1_, the protein dimer *S*_2_, the unoccupied promoter site *S*_3_, the dimer-bound promoter site *S*_4_ and the mRNA *S*_5_. These species participate in nine reactions, which are recorded in Table 2, along with the associated propensities and kinetic parameter values. The system is analyzed on the time interval [0, 2] with the initial conditions *X*(0) = [10, 20, 10, 40, 0]. For these parameter values the reversible reactions, (*R*_1_, *R*_2_), and (*R*_3_, *R*_4_,) are fast and the others are slow, thus the model is stiff [27].

**Table 2:** Genetic positive feedback loop model.

|  | Reaction | Propensity | Parameter value |
| --- | --- | --- | --- |
| $R_1$ | $S_1 + S_1 \xrightarrow{c_1} S_2$ | $a_1 = c_1 X_1(X_1 - 1)/2$ | 5000 |
| $R_2$ | $S_2 \xrightarrow{c_2} S_1 + S_1$ | $a_2 = c_2 X_2$ | $10^6$ |
| $R_3$ | $S_2 + S_3 \xrightarrow{c_3} S_4$ | $a_3 = c_3 X_2 X_3$ | 5000 |
| $R_4$ | $S_4 \xrightarrow{c_4} S_2 + S_3$ | $a_4 = c_4 X_4$ | $10^6$ |
| $R_5$ | $S_3 \xrightarrow{c_5} S_3 + S_5$ | $a_5 = c_5 X_3$ | 10 |
| $R_6$ | $S_4 \xrightarrow{c_6} S_4 + S_5$ | $a_6 = c_6 X_4$ | 20 |
| $R_7$ | $S_5 \xrightarrow{c_7} S_5 + S_1$ | $a_7 = c_7 X_5$ | 1 |
| $R_8$ | $S_1 \xrightarrow{c_8} \emptyset$ | $a_8 = c_8 X_1$ | 0.8 |
| $R_9$ | $S_5 \xrightarrow{c_9} \emptyset$ | $a_9 = c_9 X_5$ | 7 |

The CIT-2 and CFD-2 methods were applied to approximate *the second-order sensitivities with respect to parameters c*_7_ *and c*_8_ by finite-differences. A relative perturbation parameter *h* = 0.02 (around 2% of the values of the perturbed parameters) was employed. We present below the results obtained from 100, 000 quadruples of sample paths generated with two finite-difference methods for estimating 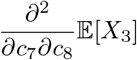. The CIT-2 technique used the tolerance value *ε* = 0.03. First, we assessed the performance of the two stochastic simulation strategies on which these estimators are based, the variable step size implicit tau-leaping method (for CIT-2) and the NRM (for CFD-2), for the nominal values of the parameters. Figure 3(a) and Figure 3(b) show the evolutions of the mean and standard deviation of the molecular amounts of *S*_3_ for both methods, respectively. The two methods yield almost identical approximations of the means, which validates the accuracy of the adaptive implicit *τ* -leaping technique. The CIT-2 and CFD-2 estimates of the second-order sensitivities of the molecular counts of *S*_3_ over time are depicted in Figure 3(c). The two estimators produced comparable results, with the CIT-2 algorithm generating a smoother curve than the CFD-2. The standard deviations of the two sensitivity estimators for the molecular counts of species *S*_3_, displayed in Figure 3(d), are comparable over the time-interval, as illustrated by the results in Figure 3(c). As before, the gain in *computational efficiency* of the CIT-2 over the CFD-2 method is substantial

**Figure 3:**
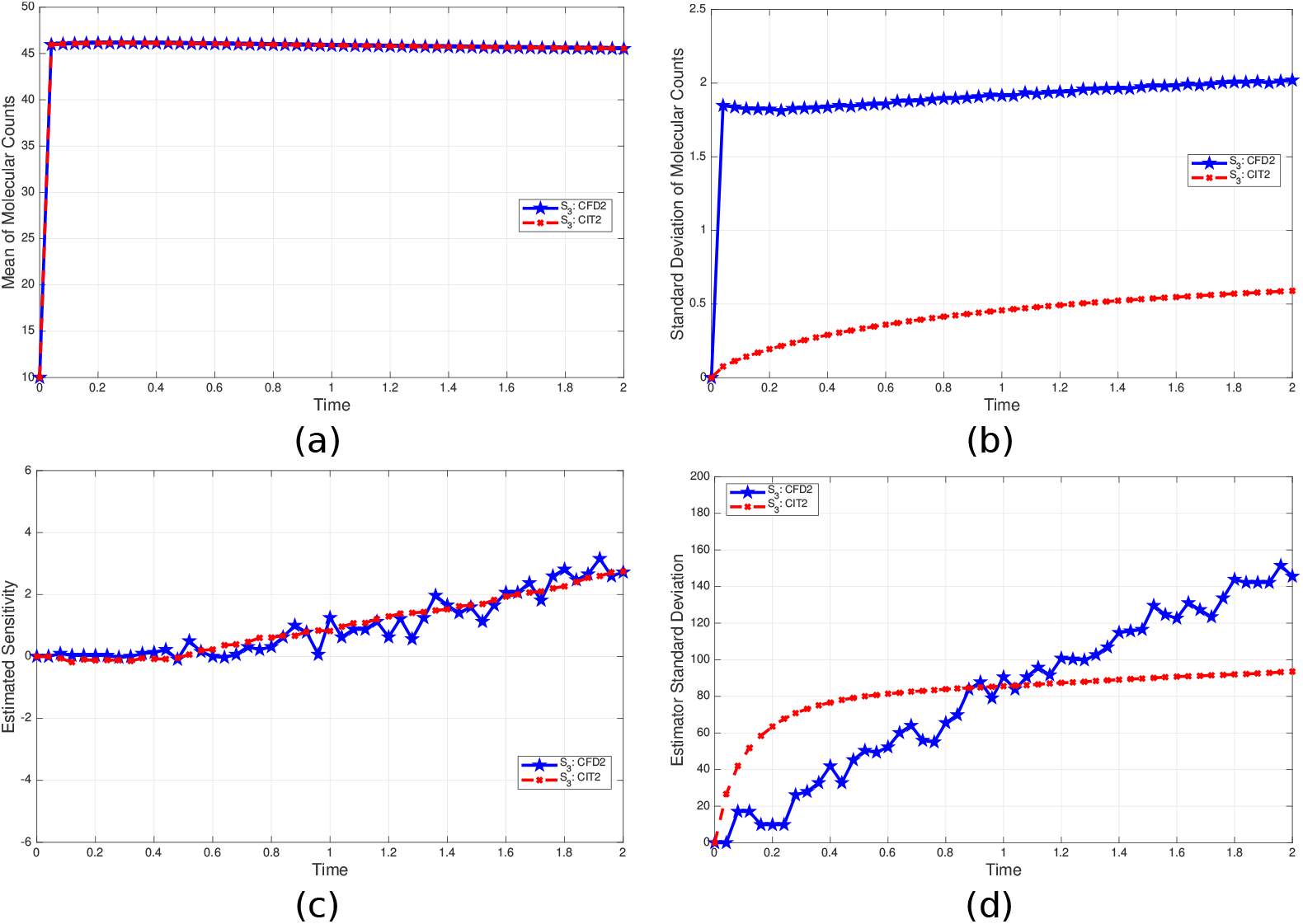
Genetic positive feedback loop model: (a) Mean and (b) Standard deviation of the molecular numbers of *S*_3_ using the variable timestep implicit tau-leaping scheme and the NRM at the nominal values of the parameters. (c) Second-order sensitivity of the molecular numbers of *S*_3_ with respect to parameters *c*_7_ and *c*_8_ with the CFD-2 and CIT-2 schemes (d) Standard deviation of the above sensitivity estimators. 100,000 quadruples of trajectories were generated for each scheme.

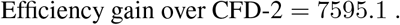

The CIT-2 method is several orders of magnitude faster than the CFD-2 scheme for estimating second-order sensitivities on this stiff biochemical model of moderate size, for the set of parameter values utilized. The two techniques have comparable accuracy.

### 4.3 Mutually Inhibiting Enzymes Model

The final biochemical model represents a mutually inhibiting enzymes system [19, 25, 39], which also appeared with a spatial distribution in [26]. In this model, eight species take part in twelve reactions. The reaction channels are described in Table 3, which also includes the associated propensities and kinetic parameter values. The initial molecular amounts for the eight species are given in Table 4. This system, studied on the interval [0, 10], exhibits stiffness.

**Table 3:** Mutually inhibiting enzymes system.

|  | Reaction | Propensity | Parameter value |
| --- | --- | --- | --- |
| $R_1$ | $E_A \xrightarrow{c_1} E_A + A$ | $a_1 = c_1[E_A]$ | 150 |
| $R_2$ | $E_B \xrightarrow{c_2} E_B + B$ | $a_2 = c_2[E_B]$ | 150 |
| $R_3$ | $E_A + B \xrightarrow{c_3} E_AB$ | $a_3 = c_3[E_A][B]$ | 0.0001 |
| $R_4$ | $E_AB \xrightarrow{c_4} E_A + B$ | $a_4 = c_4[E_AB]$ | 0.6 |
| $R_5$ | $E_AB + B \xrightarrow{c_5} E_AB_2$ | $a_5 = c_5[E_AB][B]$ | 0.0001 |
| $R_6$ | $E_AB_2 \xrightarrow{c_6} E_AB + B$ | $a_6 = c_6[E_AB_2]$ | 0.6 |
| $R_7$ | $A \xrightarrow{c_7} \emptyset$ | $a_7 = c_7[A]$ | 0.5 |
| $R_8$ | $E_B + A \xrightarrow{c_8} E_BA$ | $a_8 = c_8[E_B][A]$ | 0.0001 |
| $R_9$ | $E_BA \xrightarrow{c_9} E_B + A$ | $a_9 = c_9[E_BA]$ | 0.6 |
| $R_{10}$ | $E_BA + A \xrightarrow{c_{10}} E_BA_2$ | $a_{10} = c_{10}[E_BA][A]$ | 0.0001 |
| $R_{11}$ | $E_BA_2 \xrightarrow{c_{11}} E_BA + A$ | $a_{11} = c_{11}[E_BA_2]$ | 0.6 |
| $R_{12}$ | $B \xrightarrow{c_{12}} \emptyset$ | $a_{12} = c_{12}[B]$ | 0.5 |

**Table 4:** List of initial molecular amounts for the mutually inhibiting enzymes system.

|  | Species | Initial value |
| --- | --- | --- |
| 1. | $A$ | 2000 |
| 2. | $B$ | 1500 |
| 3. | $E_A$ | 950 |
| 4. | $E_B$ | 950 |
| 5. | $E_AB$ | 200 |
| 6. | $E_AB_2$ | 50 |
| 7. | $E_BA$ | 200 |
| 8. | $E_BA_2$ | 50 |

For this model, we considered the *second-order sensitivities with respect to parameters c*_1_ *and c*_2_ and we generated 30,000 quadruple of trajectories with the CDF-2 and CIT-2 schemes to approximate these sensitivities with a relative perturbation *h* = 0.75 which represents 0.5% of the value of the perturbed parameters *c*_1_ and *c*_2_. The CIT-2 scheme was applied with the tolerance value *ε* = 0.03. The plots of the time evolutions of the mean and standard deviations of the *E*_*B*_*A*_2_ molecular counts at the nominal values of the parameters, with the adaptive implicit *τ* -leaping technique (for CIT-2) and the NRM (for CFD-2), are displayed in Figure 4(a) and Figure 4(b), respectively. Remark that the means are in excellent agreement and so are the standard deviations, which indicate that the adaptive tau-leaping scheme with the selected value of the tolerance *ε* achieves the same accuracy as the exact next reaction method on which the CFD-2 strategy is built.

**Figure 4:**
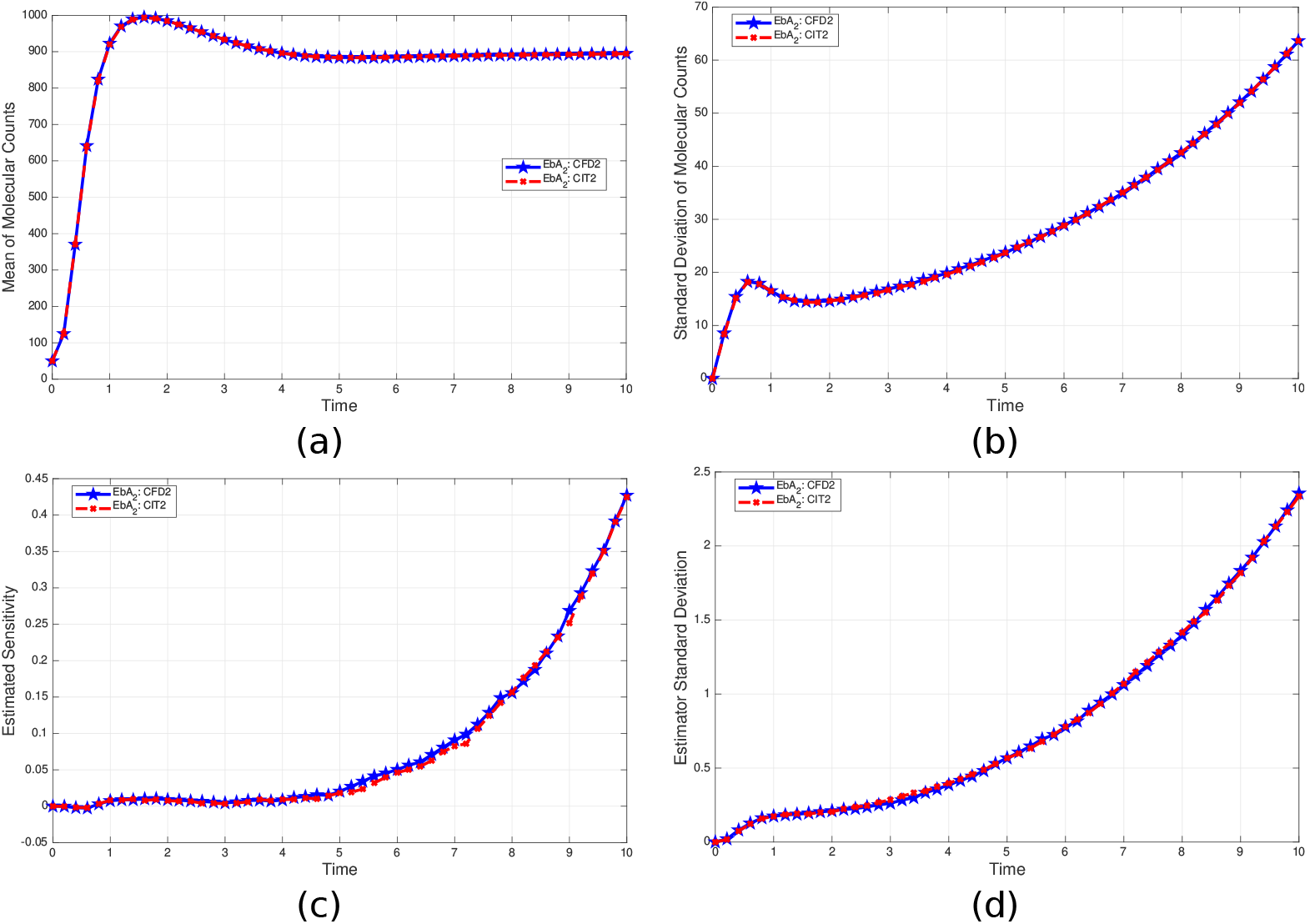
Mutually inhibiting enzymes model: (a) Mean and (b) Standard deviation of the molecular abundance of the *E*_*B*_*A*_2_ species with the adaptive implicit *τ* -leaping and the NRM schemes. (c) CFD-2 and CIT-2 sensitivity estimators of second-order with respect to *c*_1_ and *c*_2_ of the molecular counts of *E*_*B*_*A*_2_. (d) Standard deviation of the above sensitivity estimators. 30,000 quadruples of sample paths were simulated.

Figure 4(c) shows the estimated second-order sensitivity 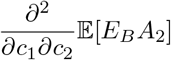 as function of time, produced by the CFD-2 and CIT-2 methods. The curves show nearly identical results. In addition, the standard deviation of the two sensitivity estimates against time are presented in Figure 4(d). It can be observed that the two strategies yield the same variance of the estimator. The close match of the two sensitivity approximations validates the accuracy of the proposed CIT-2 method, as evident from Figure 4(d).

Finally, the computational efficiency of the CIT-2 relative to the CFD-2 method for this particular simulation is determined to be

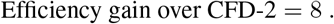

This shows that the proposed method achieves almost an order of magnitude reduction in computational cost, on the stiff model analyzed, which has many reactions and biochemical species.

## 5 Conclusions

Second-order sensitivity analysis of biochemical reaction models is a fundamental tool for parameter estimation, model calibration, and experimental design. In this work, we have proposed a novel coupled implicit tau-leaping method for second-order parametric sensitivity analysis of stochastic discrete models of biochemical networks, referred to as CIT-2. The CIT-2 method is a finite-difference technique for estimating the entries of the Hessian, the second-order sensitivities with respect to model parameters, of biochemical systems represented by the Chemical Master Equation model. This strategy constructs quadruples of sample paths for a finite difference approximation, a nominal path, two once perturbed paths and one doubly-perturbed path, utilizing an adaptive implicit tau-leaping strategy on a common sequence of time-steps. It is based on a tight coupling of the associated stochastic processes, which shared Poisson random variables with the intensities akin to those of the coupled finite-difference method of second-order (CFD-2) introduced in [42]. While the underlying strategy of the CFD-2 scheme is an exact stochastic simulation method, our technique uses the approximate implicit tau-leaping technique together with a variable times-step selection adapted from that of Cao et al. [7]. The implicit tau-leaping approach blended the variance reduction of a strong coupling of the nominal and perturbed processes with numerical stability to effectively deal with the computational challenges posed by stiffness. Stiffness is exhibited in numerous realistic stochastic models of biochemical systems. Our approach makes second-order sensitivity analysis feasible for such systems, for which exact simulation strategies and explicit *τ* -leaping schemes are computationally impractical.

The advantages of the proposed method were illustrated on three stiff biochemical models: the decay-dimerization, a genetic positive feedback loop system and a mutually inhibiting enzymes network, which are models of small to moderate sizes. For these models, the CIT-2 and CFD-2 sensitivity estimates are in very good agreement, for a similar level of accuracy. The numerical experiments show that the variance of the CIT-2 estimator is less or comparable to that of the existing estimator. Moreover, the proposed method requires *substantially less computational effort* compared to the published technique on the models tested, often of several orders of magnitude. This gain in efficiency depends on the model, and is expected to grow as the degree of stiffness increases. Our technique is expected to outperform existing strategies for models which are stiff to very stiff. It may be employed with central finite-difference schemes and it may be utilized for estimating sensitivities of higher moments of the system output. We intend to extend the approach introduced in this paper to higher-order sensitivity estimations.

## Acknowledgements

This research was supported by a grant from the National Sciences and Engineering Research Council of Canada (NSERC) - Grant No. RGPIN-2020-05469, Toronto Metropolitan University Mathematics Department and Faculty of Science Dean’s Research Fund.

